# Daily Acute Intermittent Hypoxia Elicits Age & Sex-Dependent Changes in Molecules Regulating Phrenic Motor Plasticity

**DOI:** 10.1101/2024.05.07.592914

**Authors:** Jayakrishnan Nair, Alexendria B. Marciante, Carter Lurk, Mia N. Kelly, Gordon S. Mitchell

## Abstract

Acute intermittent hypoxia (AIH) elicits a form of respiratory motor plasticity known as phrenic long-term facilitation (LTF). Repetitive daily AIH (dAIH) exposure enhances phrenic LTF, demonstrating a form of metaplasticity. Two additional factors impacting phrenic LTF are age and sex. For example, moderate AIH-induced phrenic LTF decreases with age in males, but increases in middle-aged females. However, little is known concerning cellular mechanisms of dAIH effects or age-dependent sexual dimorphism in phrenic LTF. Moderate AIH elicits distinct signaling cascades within phrenic motor neurons initiated by 5HT2 receptors (Q pathway) *versus* adenosine 2A or 5HT7 receptors (S pathway), respectively. The Q and S pathways interact *via* mutual crosstalk inhibition, a powerful regulator of phrenic LTF. To test the hypothesis that dAIH, age and sex effects on phrenic LTF are associated with differential expression of molecules known to regulate the Q and S pathways, we assessed mRNA of key regulatory molecules in ventral cervical homogenates from spinal segments containing the phrenic motor nucleus from young (3 month) and middle-aged (12 month) male and female Sprague-Dawley rats. Since CNS estrogen levels impact molecules regulating the Q and/or S pathways, mRNA was correlated with serum estradiol. Rats (n=8/group) were exposed to sham (21% O_2_) or dAIH (15, 1 min episodes of 10.5% inspired O_2_ per day) for 14 days, and sacrificed 24 hours post-dAIH. mRNA for molecules known to regulate phrenic LTF were assessed *via* RT PCR, including: brain derived neurotrophic factor (*Bdnf*); serotonin 2A (*Htr2a*); 2B (*Htr2b*); and (*Htr7*) receptors; adenosine 2a (*Adora2a*) receptors; exchange protein activated by cAMP (*Epac1*); p38 MAP kinase [*Mapk14* (α) & *Mapk11* (β)]; PKA catalytic subunit (*Prkaa1*); PKA regulatory subunit (*Prkar1a*); fractalkine (*Cx3cl1*); phosphodiesterase type 4 (*Pde4b*); NAPDH–gp91 (*Cybb*) and p47 (*ncf1*); and the PKCδ isoform (*Prkcd*). Significantly higher *Pde4b*, *Adora2a*, and *Prkcd* mRNA were found in young and middle-aged females *versus* age-matched males; *Epac1* was elevated, but only in young females (p<0.001). *Ncf1* was increased in middle-aged versus young adult rats of both sexes (p<0.01). *Ncf1, Cx3cl1*, *Adora2a* and *Prkcd* mRNA were reduced by dAIH in middle-aged females (p<0.01), but not other groups. Serum estradiol levels positively correlated with *Epac1* (r^2^=0.29, p=0.002), *Mapk14* (r^2^=0.31, p=0.001), *Mapk11* (r^2^=0.20, p=0.014), and *Prkar1a* (r^2^=0.20 p=0.013) mRNA. With higher serum estradiol levels, dAIH decreased *Mapk14* mRNA (slope difference p=0.001). Thus, age, sex and dAIH preconditioning influence molecules known to regulate the Q and S pathways to phrenic motor facilitation. These novel findings advance our understanding of phrenic LTF, and inform translational research concerning the therapeutic potential of dAIH to treat breathing deficits in individuals of different ages or sex.

## INTRODUCTION

Acute intermittent hypoxia (AIH) persistently increases phrenic nerve activity, an effect known as phrenic long-term facilitation (pLTF). The magnitude of AIH-induced pLTF is enhanced by pre-conditioning with daily AIH (dAIH, (Fields & Mitchell, 2015) or repetitive AIH (3 times/week; (Fields & Mitchell, 2015; MacFarlane et al., 2018) *via* unknown mechanisms. Understanding cellular mechanisms and factors that regulate dAIH-enhanced plasticity is important since repetitive AIH is emerging as a therapeutic modality to treat life-threatening respiratory (and non-respiratory) motor deficits caused by spinal cord injury and other neuromuscular disorders (Vose et al., 2022). Based on our nuanced understanding of cellular mechanisms giving rise to pLTF (Mitchell & Baker, 2022), we selected key molecules known to regulate pLTF and investigated dAIH effects on their (mRNA) gene expression in ventral cervical homogenates containing the phrenic motor nucleus. Further, we investigated the impact of age and sex on these changes.

Activation of key phrenic motor neuron Gq (Q pathway) or Gs (S pathway) protein–coupled metabotropic receptors and subsequent intracellular signaling cascades guided the present study of dAIH-induced gene expression (Dale-Nagle et al., 2010; Mitchell & Baker, 2022). With moderate AIH (PaO_2_ ≳ 40 mmHg), Gq-coupled serotonin (5-HT2) receptor-dependent pLTF occurs (Baker-Herman & Mitchell, 2002; Fuller et al., 2001; Kinkead et al., 1998; MacFarlane et al., 2008). Q pathway-driven plasticity requires downstream ERK MAP kinase, TrkB, and protein kinase C-θ (PKCθ) (Dale et al., 2017; Devinney et al., 2015; Hoffman et al., 2012) activity, new BDNF protein synthesis (Baker-Herman & Mitchell, 2002), reactive oxygen species (ROS) formation (MacFarlane et al., 2009; MacFarlane et al., 2008), and neuronal nitric oxide synthase (nNOS) activity (MacFarlane et al., 2014). Conversely, with severe AIH (PaO_2_ ≲ 30 mmHg), pLTF is driven by GS-coupled adenosine 2A (A2a) and 5-HT7 receptor activation (Golder et al., 2008; Hoffman & Mitchell, 2011; Nichols et al., 2012; Nichols & Mitchell, 2021). S pathway signaling requires synthesis of an immature TrkB isoform (*versus* BDNF), PI3 kinase/Akt signaling (*versus* ERK) (Golder et al., 2008). However, with PaO_2_ between ∼30-35 mmHg, Q-S pathway co-activation cancels pLTF due to crosstalk inhibition (**Figure 1**) (Perim & Mitchell, 2019).

**Figure 1:**
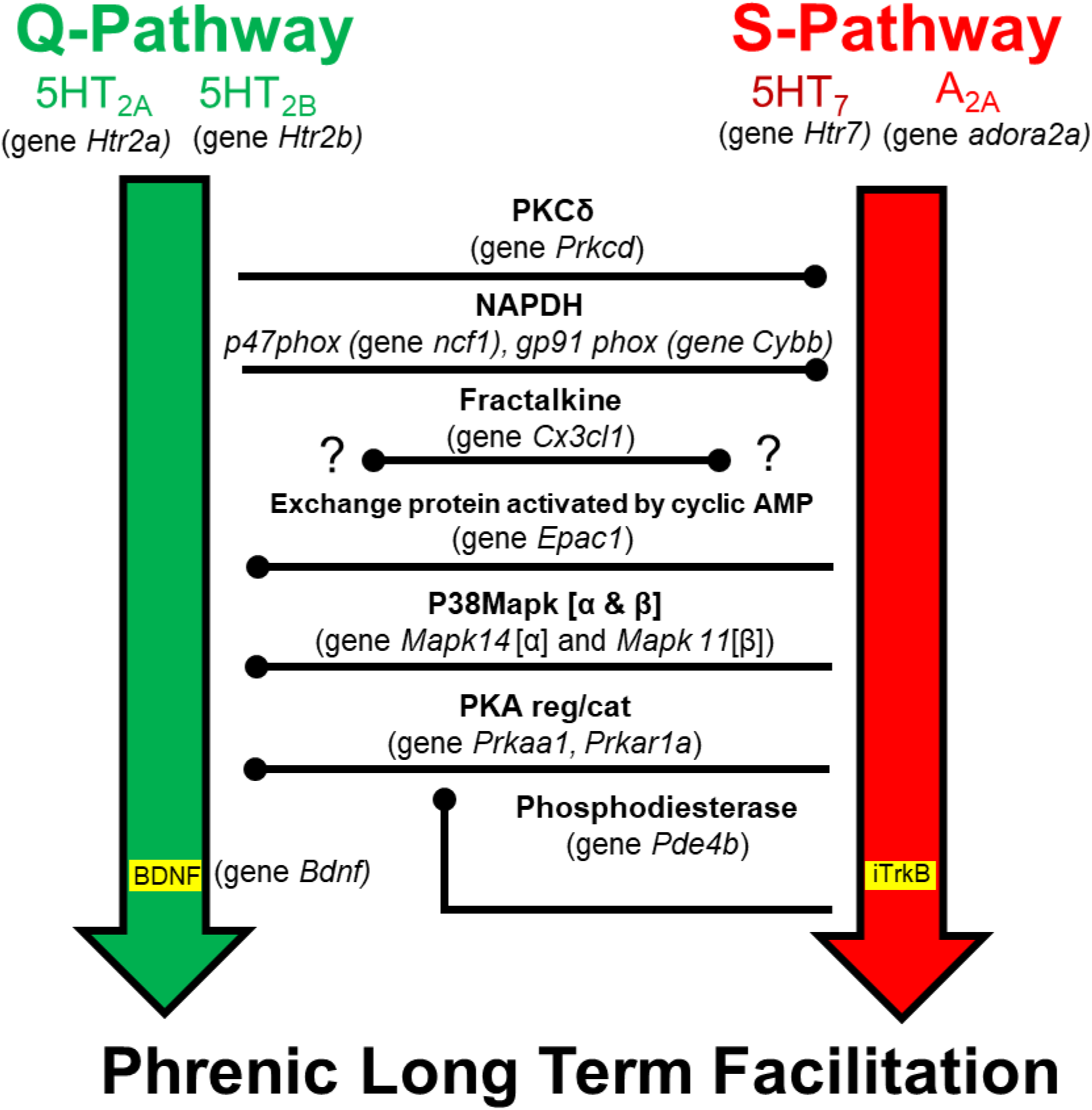
Conceptual framework guiding investigation of intra-ceullular signaling cascades driving or regulating AIH induced phrenic long-term facilitation. Depicted are the Serotonin−5HT2A & 5HT2B, BDNF dependent (Q pathway; green) and the 5HT7 & Adenosine 2A, immature TrKB (iTrkB) dependent (S pathway; red). Q to S pathway cross talk inhibition is mediated by PKCδ or NADPH (p47 and gp91 are key subunit genes). S to Q pathway cross talk inhibition is mediated by Protein Kinase A (PKA; key subunit genes encode regulatory and catalytic domains), with suspected involvement of p38Map kinase. Phosphodiesterase indirectly inhibits PKA activity by decreasing cAMP levels, liberating the Q pathway and suppressing S pathway activation. Fractalkine protein activates its receptor on nearby microglia, triggering them to increase extracellular adenosine, thus suppressing the Q pathway and favoring the S pathway. Genes associated with each of these molecules are shown in parenthesis.

AIH protocols consisting of shorter hypoxic episodes minimize hypoxia-evoked spinal adenosine accumulation and S-Q crosstalk inhibition (Marciante et al., 2023). However, multiple conditions can shift the serotonin/adenosine balance (Mitchell & Baker, 2022). For example, with an injured spinal cord, heightened tissue hypoxia from systemic hypotension and/or pericyte capillary constriction (Li et al., 2017) may increase Q-S crosstalk inhibition, even at the same PaO_2_ (Perim, Gonzalez-Rothi, et al., 2021; Perim et al., 2023). Time of the day (Marciante et al., 2023), age and sex (Behan et al., 2002; Behan et al., 2003; Zabka et al., 2001a, 2001b) also regulate pLTF due to changes in Q-S cross talk inhibition (Marciante & Mitchell, 2023; Marciante et al., 2023). Phrenic LTF elicited by AIH consisting of 1 minute hypoxic episodes is attenuated in the rodent active (*versus* rest) phase due to diurnal variations in spinal adenosine levels (Marciante et al., 2023). Phrenic LTF (rest phase) decreases with age in males, but increases in middle-aged females, with a pronounced estrus cycle variation; some of these effects may arise from estradiol effects on the Q & S pathways or their cross-talk inhibition (Dougherty et al., 2017; Zabka et al., 2006).

Pretreatment with daily AIH (dAIH) enhances phrenic LTF—a form of metaplasticity (Fields & Mitchell, 2015; Mitchell & Johnson, 2003). Although mechanisms whereby dAIH enhances phrenic LTF are unknown, they may relate to: 1) Q pathway amplification (MacFarlane et al., 2018; Satriotomo et al., 2012); or 2) minimizing S-Q crosstalk constraints (Hoffman et al., 2010; Mitchell & Baker, 2022). Since aging increases Q-to-S cross talk inhibition due to increased spinal adenosine in geriatric male and female rats (Marciante & Mitchell, 2023), differential age and sex effects of dAIH must be considered. Since pLTF is impacted greatly by the estrus cycle in females, it is important to account for estrogen effects on the Q and S pathways (Dougherty et al., 2017; Zabka et al., 2006). Here, we begin exploring these relationships by screening mRNA for molecules known to regulate pLTF in ventral cervical homogenates with and without dAIH in young and middle-aged male and female rats.

We hypothesize that 2 weeks of dAIH: 1) decreases ventral cervical spinal mRNA of molecules mediating Q and S pathway crosstalk inhibition; and/or 2) increases mRNA of key Q pathway molecules. We further hypothesize that age and sex regulate mRNA of molecules regulating phrenic LTF, and that these effects correlate with serum estradiol levels.

## MATERIALS AND METHODS

### Animals

All experimental procedures were approved by the Institutional Animal Care and Use Committee at the University of Florida and conformed to policies detailed in the National Institutes of Health Guide for the Care and Use of Laboratory Animals. Experiments were performed on naïve 3-month-old male (297 ± 16 g, n=8) and female (243 ± 7 g, n=8), and on naïve 12-month-old, middle-aged, male (493 ± 16 g, n=8) and female (286 ± 55 g, n=8) Sprague Dawley rats (208A Colony, Inotive formally known as Envigo, IN). Rats were housed in pairs in individually ventilated cages and maintained in a mixed-sex rodent room within an AAALAC-accredited animal facility for at least one week prior to use. The ambient conditions at the facilities were maintained at 24°C with 12 hr light-dark cycles (6 am-6 pm) with access to food and water *ad libitum*. Sample size estimation was based on prior studies (McIntosh & Dougherty, 2019; Zabka et al., 2001b).

### Daily AIH protocol

Rats were randomly assigned to receive either daily poikilocapnic AIH (dAIH) or continuous normoxia (Sham exposures consisting of 15, 1-min episodes of hypoxia (10.5% inspired O_2_) separated by 1-min intervals of normoxia; AIH was repeated daily for 14 consecutive days (**Figure 2**). On exposure days, the rats were placed in individual, custom-designed cylindrical exposure chambers with flat inserts to support the animals and enable urine to pass below in the platform. Each chamber was connected to a computer-controlled programmable mass flow controller system (Flow Commander, Therapeutiq Research, Kansas, USA) to regulate chamber gas flow (∼6 L/min) and gas composition, with targeted inspired O_2_ levels (Fig. 1). Cotton gauze placed close to the airflow inlet diffused the incurrent gas flow. This system allowed automated cycling between hypoxia and room air or sham normoxic exposures. Rats were allowed ∼30 min to acclimate to the chamber before AIH began. Exposures were given between 8:30-11 AM each day, and then the rats were returned to their home cages.

**Figure 2:**
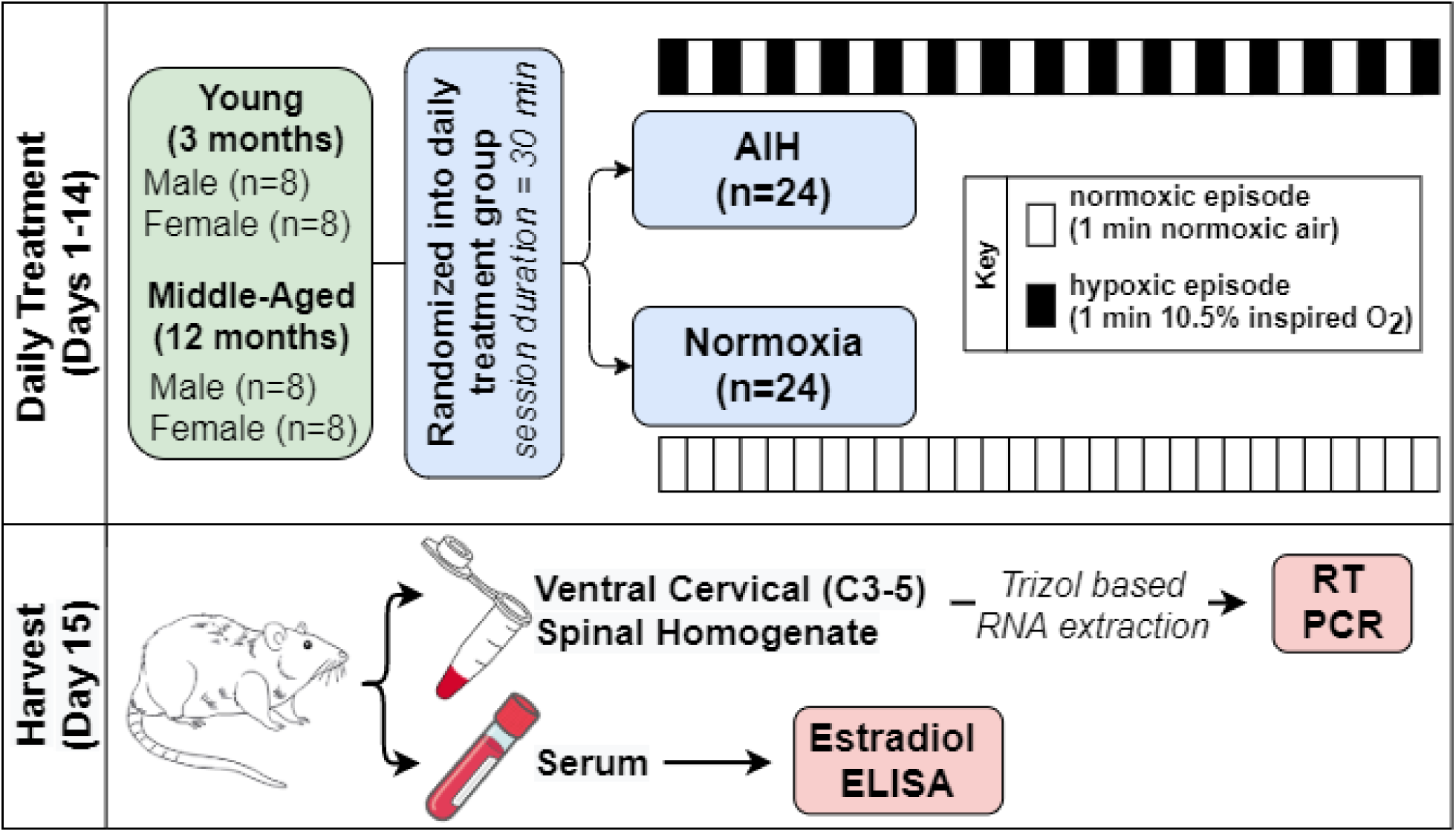
Schematic of the experimental design. Young (3 months) and middle aged (12 month) rats of both sexes were exposed to daily AIH for 14 consecutive days (15, 1 minute hypoxic episodes with 1 ½ min normoxic intervals) or Sham (normoxia). 24 hours after the last exposure, rats were sacrificed under deep isoflurane anesthesia and ventral cervical spinal cord tissues were dissected and flash frozen for mRNA analysis using RT PCR. At the same time serum samples from these rats were collected and stored for estradiol ELISA.

### Necropsy and sample collection

24 hrs after dAIH, rats were euthanized under deep isoflurane anesthesia (4%) and decapitated *via* rodent guillotine. Prior to sacrifice, the estrus cycle of female rats was determined via vaginal swabs as described by Marcondes et al,.(Marcondes et al., 2002) Blood was collected in a serum separator vial and allowed to clot for 20-30 min at room temperature. The sample was centrifuged at 3000RPM for 20 minutes. Separated serum was removed and stored at −80°C for ELISA-based estradiol analysis. Following decapitation the C3-C5 spinal segment was rapidly removed and immersed in ice-cold, sterile phosphate buffer solution (PBS). C3-C5 spinal cord samples were transferred to a freezing microtome (−22°C) and cut at the level of the central canal with a double edge blade and saved in RNA later stabilization solution (Thermo Fisher Scientific); these maneuvers were achieved within 14±1.1 min from the time of sacrifice. Ventral portions of C3-C5 were placed in RNAlater stabilization solution and stored at 4° C overnight before transferring to −80°C for long-term storage. On the RNA isolation day, frozen samples were thawed on ice and wet weights were recorded (38±5.6 g; mean ± SD).

### RNA isolation and quantification procedures

Tissue samples were homogenized in 800µl TRIzol Reagent (Invitrogen) using Bead Mill 4 Homogenizer (Thermo Fisher Scientific). The extracted product was cleaned by ethanol precipitation and transferred to the RNeasy spin column (RNeasy kit, Qiagen), and purified. RNA concentration was quantified via spectrophotometry (ʎ= 260 nm; NanoDrop model 2000C, ThermoFisher Scientific). RNA purity was estimated by the absorbance ratio A_₂₆₀_/A_₂₈₀_ (all samples had a ratio from 1.8 to 2.0).

### Real-time reverse transcription-polymerase chain reaction

Extraction and quantification procedures as described by Kelly et al.(Kelly et al., 2020) were used for RT-PCR. Briefly, the first strand complementary DNA (cDNA) was synthesized from 2.5µg of total RNA using random primers in SuperScript VILO cDNA Synthesis Kit (Invitrogen). The resulting cDNAs were diluted to 4 ng/µL and used as templates in real-time quantitative polymerase chain reactions (PCRs; QuantStudio3; Applied Biosystems). TaqMan oligonucleotide primers and probe sets (TaqMan Gene Expression Assays; Applied Biosystems) used for PCR are listed in Table 1. Twenty µL reactions were prepared. Reaction mixtures consisted of 5 µL cDNA (4 ng/µL), 10 µL TaqMan Fast Advanced Master Mix (Applied Biosystems), 4 µL DEPC-treated RNAse-free water, and 1 µL of the corresponding TaqMan gene expression assay. Taq-Man primers and probes were used in duplicate reactions, with the same cycling conditions (50°C for 2 min, and 95°C for 2 min, followed by 40 cycles of 95°C for 1 s and 60°C for 20 s). Negative control reactions confirmed minimal contamination. mRNA in ventral C3–C5 homogenates were normalized as a difference from the amplification cycle at the 18s mRNA threshold. Quantification was performed *via* the ΔΔCT method (Livak & Schmittgen, 2001); ΔCT values were calculated as CT (target gene) - CT (18s). mRNA levels are presented as fold change (2^−ΔΔCT^) from young males exposed to sham normoxia.

**Table 1:** TaqMan oligonucleotide primers and probe sets used for PCR.

| Gene Symbol | Protein Encoded | TaqMan Assay ID | Amplicon Length |
| --- | --- | --- | --- |
| *Mapk14* | p38MapKinase $\alpha$ subunit | Rn00578842_m1 | 87 |
| *Mapk11* | p38MapKinase $\beta$ subunit | Rn01407663_g1 | 65 |
| *Bdnf* | BDNF | Rn02531967_s1 | 142 |
| *Htr2a* | 5-HT2A | Rn00568473_m1 | 71 |
| *Htr2b* | 5-HT2B | Rn00691836_m1 | 81 |
| *Htr7* | 5-HT7 | Rn00576048_m1 | 85 |
| *Prkaa1* | PKA Regulatory Subunit | Rn00566036_m1 | 84 |
| *Prkar1a* | PKA Catalytic Subunit | Rn01402558_m1 | 69 |
| *Cybb* | NADPH-gp91phox | Rn00576710_m1 | 77 |
| *Epac1* | exchange protein directly activated by cAMP | Rn00572463_m1 | 65 |
| *Cx3cl1* | CX3CL1 Fractalkine | Rn00593186_m1 | 74 |
| *ncf1* | NADPH-p47phox | Rn00586945_m1 | 105 |
| *pde4b* | phosphodiesterase 4B | Rn00566785_m1 | 70 |
| *Prkcd* | Protein kinase C delta | Rn00440891_m1 | 58 |
| *adora2a* | Adenosine 2A | Rn00583935_m1 | 58 |
| *18s* | Eukaryotic Ribosomal RNA | Hs99999901_s1 | 187 |

### Estradiol ELISA

Frozen serum samples were thawed on ice and estradiol levels assayed *via* competitive immunoassay in rat serum (BioVendor, Cat#RTC009R) (Marciante & Mitchell, 2023). The calibration range for the assay was 2.5-1280 pg/ml. The assay was performed according to the manufacturer’s instructions; a filter-based, single-channel absorbance (range 400-750nm) microplate reader (BioTek 800 TS) was used to quantify the level of serum estradiol level. The correlation between serum estradiol and estrus cycle is presented in **Figure 3**. The estrus cycle, as determined with vaginal swab smear shows that the serum estradiol was lower in estrus (10.0 ± 2.8 pg/ml) and higher during diestrus phase (24.7 ± 2.4 pg/ml). Estradiol exhibited a high degree of variability in proestrus (15.3 ± 7.3 pg/ml; all values mean ± S.D.).

**Figure 3:**
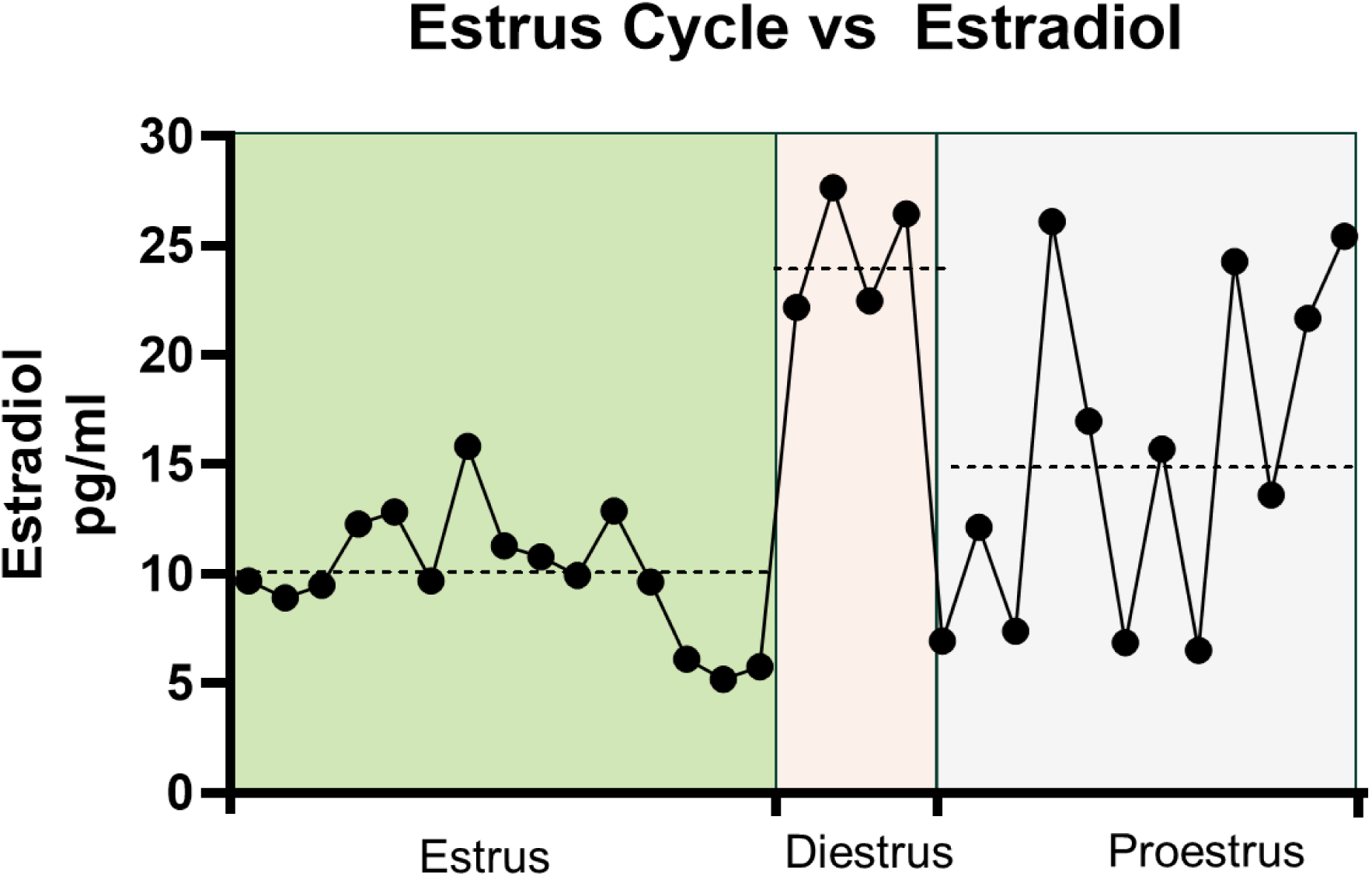
Correlation between serum estradiol and estrus cycle stage. The estrus cycle for female rats was determined *via* vaginal swab smear. The serum estrodial level was lower during estrus (m ±SD; 10.0 ± 2. 8 pg/ml) and elevated during diestrus (24.7 ± 2.4 pg/ml). Large variability characterized proestrus (15.3 ± 7.3 pg/ml). Dotted line = mean for that stage.

### Statistical analysis

To study age (3 *versus* 12 months) and sex (male *versus* female) effects on gene expression, sham rats were analyzed *via* linear regression with age and sex as predictors (SPSS). Daily AIH effects on gene expression were compared *via* pre-post paired t-tests or non-parametric Wilcoxon matched pair signed rank tests based on the Normality test with Kolmogorov-Smirnov test (Prism Graph Pad). To account for multiple comparisons, we used a conservative α error rate of p≤0.01 to determine significance for age and sex effects, as well as pre-post dAIH treatment effects. The correlation between estradiol and gene expression was assessed *via* simple linear regression (Prism Graph Pad). Significant differences in slope (α error of p≤0.05) between Sham and dAIH-exposed rats was used to determine possible effects of estradiol on dAIH-induced changes in gene expression. Outlier data points were removed from the statistical analysis if the Cook’s D was >4. Values are means ± SD.

### Elimination criterion

Each cohort had 8 rats, although data from 2 rats in the young male sham cohort were not included since the 18s reference gene values had a Cook’s D >4. Data from one middle-aged rat for Cx3cl1 and *Htr7* gene expression were removed from the analysis. Serum estradiol data from 1 young female and 2 middle-aged males could not be collected due to a sample processing error.

## RESULTS

### Effect of age-sex on expression of genes regulating plasticity

In the ventral cervical spinal cord, statistically significant effects of age-sex were detected in *Adora2a* (age: *t=2.69, p=0.012*; sex: *t=6.15, p=0.000*) and *Pde4b* (age*: t=-2.88, p=0.008*; sex: *t=3.44, p=0.002*). A statistically significant effect of age was observed with *Htr2b* (*t=4.13, p=0.001*), *Prkcd* (*t=3.01, p=0.006*), *Ncf1* (*t=4.41, p=0.001*), and *Cybb (t=2.65, p=0.013)*. Whereas statistically significant effects of sex were observed with *Cx3cl1* (*t=3.28, p=0.003*), *Epac1* (*t=5.70, p=0.001*), *Mapk14* (*t=3.94, p=0.001*), *Mapk11* (*t=3.51, p=0.002*), *Pkaa1* (*t=3.98, p=0.001*), and *Pkar1a* (*t=4.96, p=0.001*), no age or sex effects were observed in *Htr2a, Htr7* or *Bdnf mRNA levels* (Table 2).

**Table 2:**
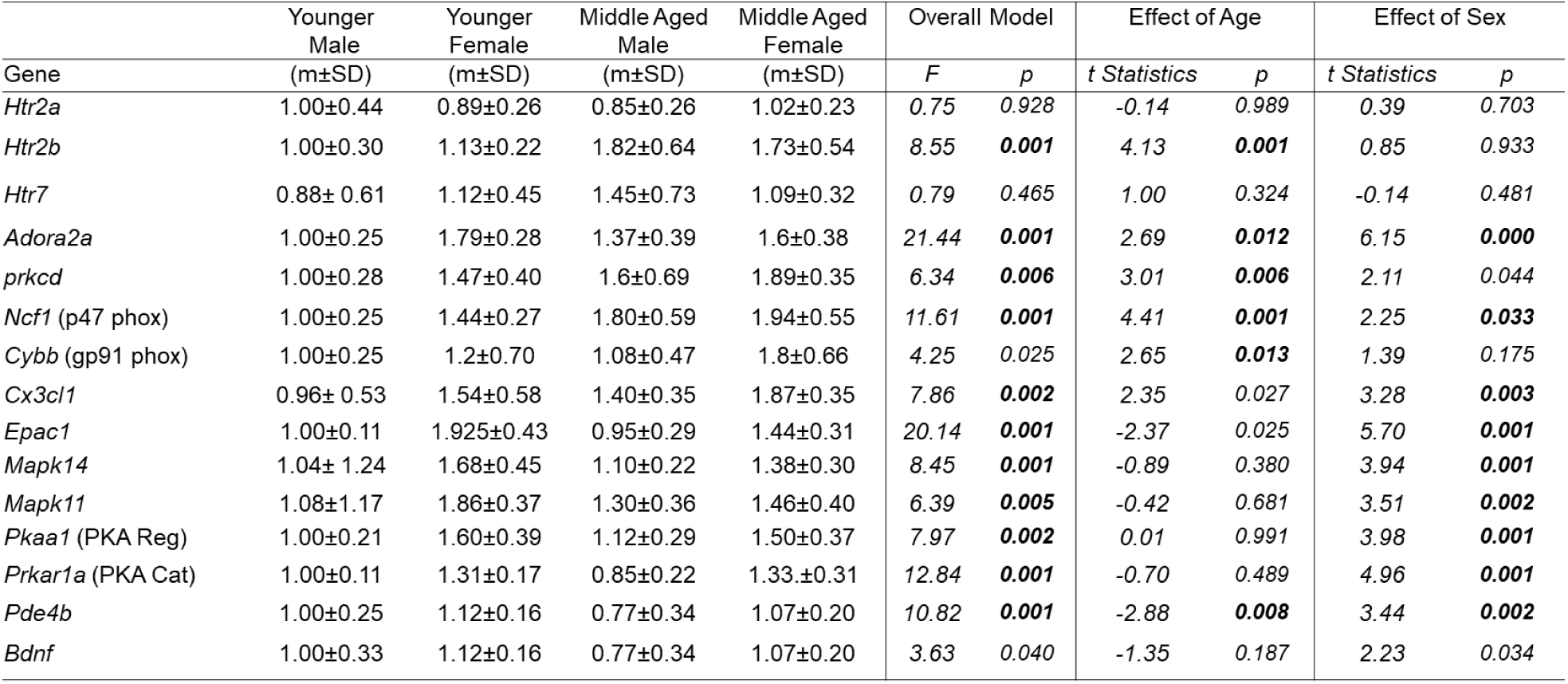
Effect of age-sex dimorphism on pro-plasticity gene expression. Significance p<0.01.

### Effect of dAIH on Q & S pathway receptor gene expression

No dAIH effects were detected on *Htr2a, Htr2b, Htr7,* or *Adora2a* receptor gene expression in younger or middle-aged males, or in younger females. Significant decreases in dAIH effects on *Htr2a* (Sham= 1.02 ±0.23; dAIH=0.78±0.14; *p=0.014*) and *Adora2a* (Sham= 1.6±0.38; dAIH=1.07±0.33; *p=0.002*) were observed in middle-aged females. However, *Htr2b* receptor expression was unaffected by dAIH in middle-aged females (Sham= 1.73±0.54; dAIH=1.16±0.33; *p=0.022*) (see **Figure 4**; Table 3).

**Figure 4:**
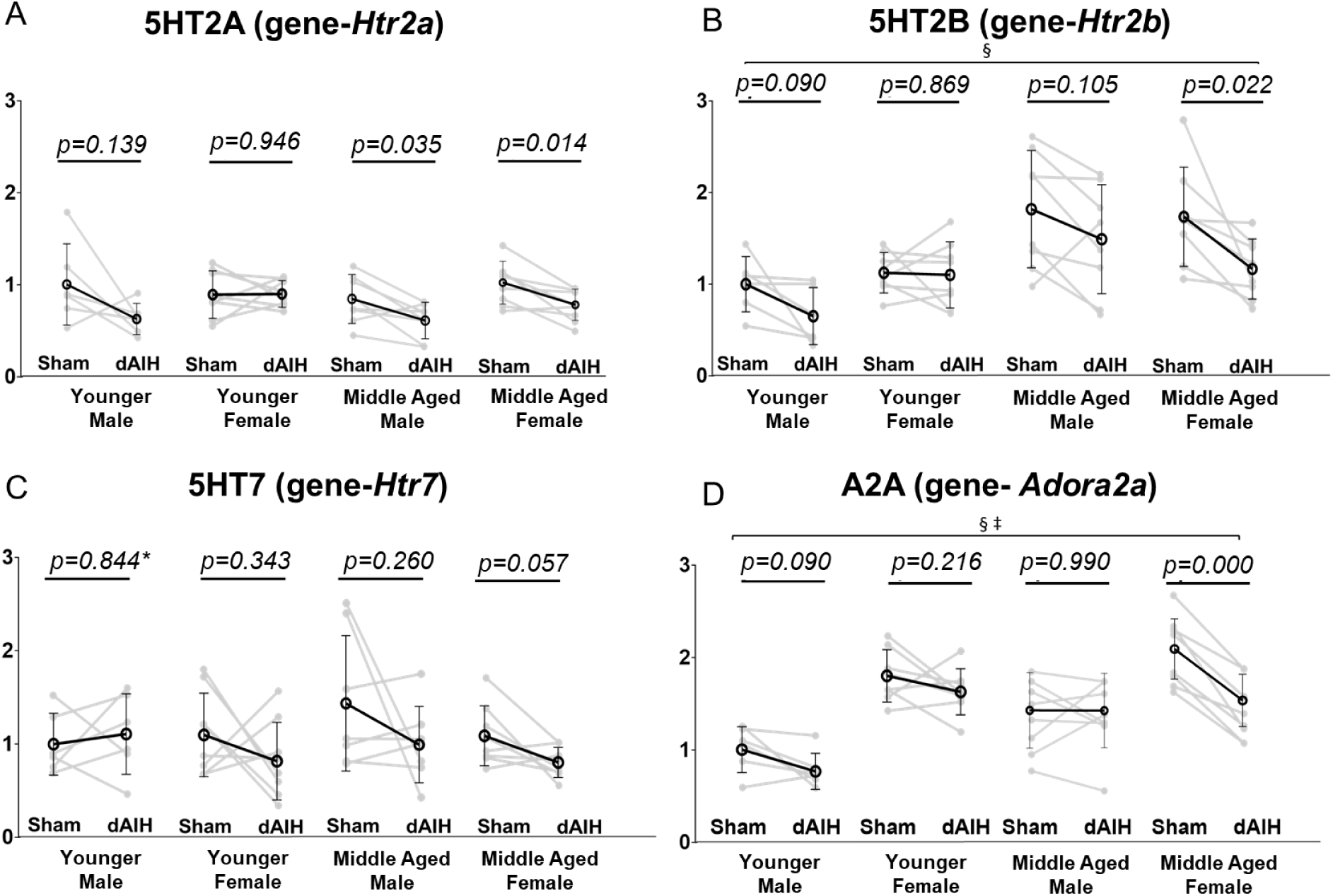
Effect of age, sex and daily AIH (dAIH) on genes associated with receptors driving the Q and S pathways: 5HT2a (*Htr2a*), 5HT2b (*Htr2b*), 5HT7 (*Htr7*), and A2a (*Adora2a*). No effect of age or sex was observed on *Htr2a* and *Htr7* expression (Panel A, C). dAIH significantly reduced *Htr2a* receptor expression in middle aged females only (Panel A). A significant effect of age was observed on *Htr2b* expression, with no significant dAIH effect in any group (Panel B). Significant age and sex effects were observed in *Adora2a* expression, with greater expression in females. dAIH significantly reduced *Adora2a* in middle aged females. §-Significant effect of age; ‡-Significant effect of sex.

**Table 3:**
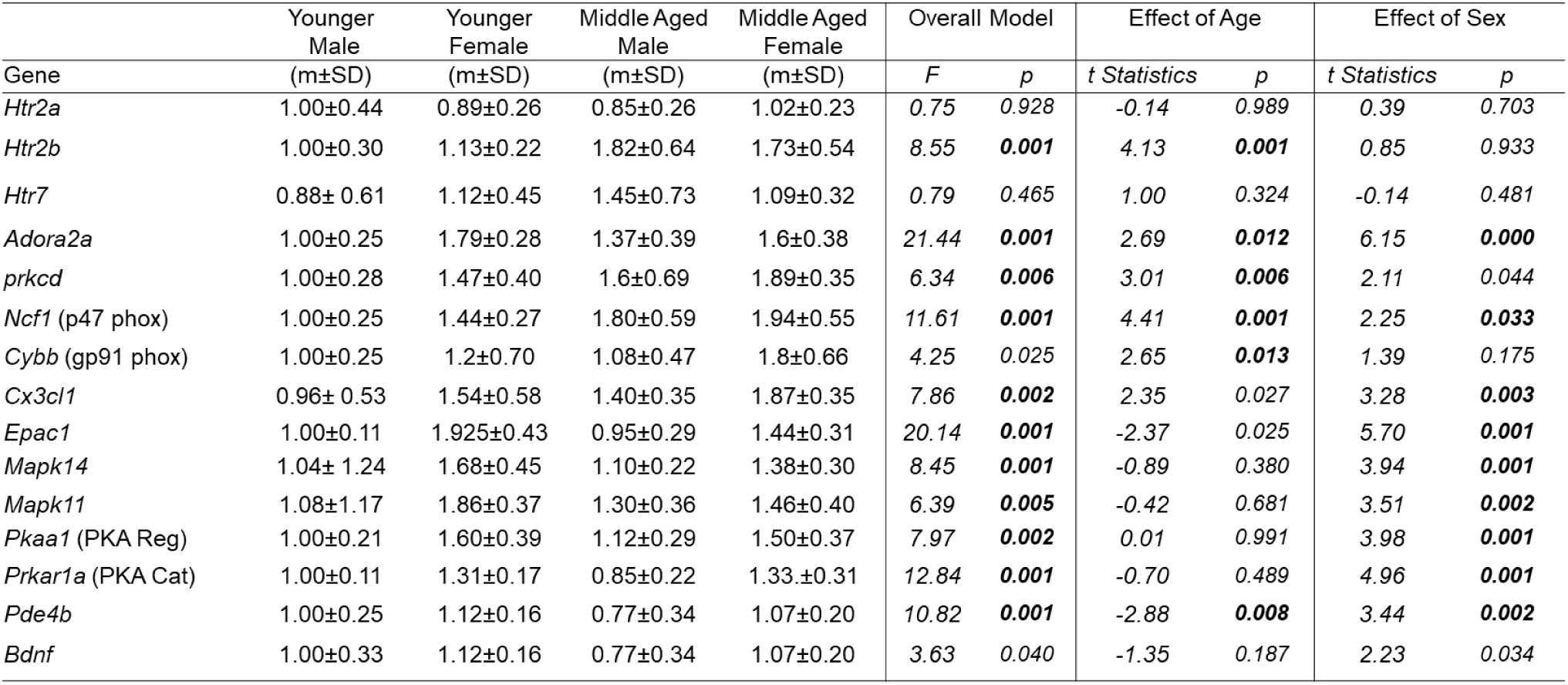
Effect of daily acute intermittent hypoxia (dAIH) on pro-plasticity gene expression. Significance p<0.01.

### dAIH effects on Q-to-S crosstalk molecule expression

No significant changes in Q-to-S cross-talk molecule gene expression were detected in younger or middle-aged males. In contrast, a significant reduction in *ncf1* was observed in younger (Sham= 1.44±0.27; dAIH= 1.12±0.13; *p=0.003*) and middle-aged females (Sham= 1.94±0.55; dAIH= 1.35±0.34; *p=0.004*). A significant reduction in *Prkcd* expression was also observed in middle-aged females (Sham= 1.89±0.35; dAIH= 0.8±0.17; *p=0.004*). No significant change in *Cybb* was detected in any dAIH-exposed group (see **Figure 5**; Table 3).

**Figure 5:**
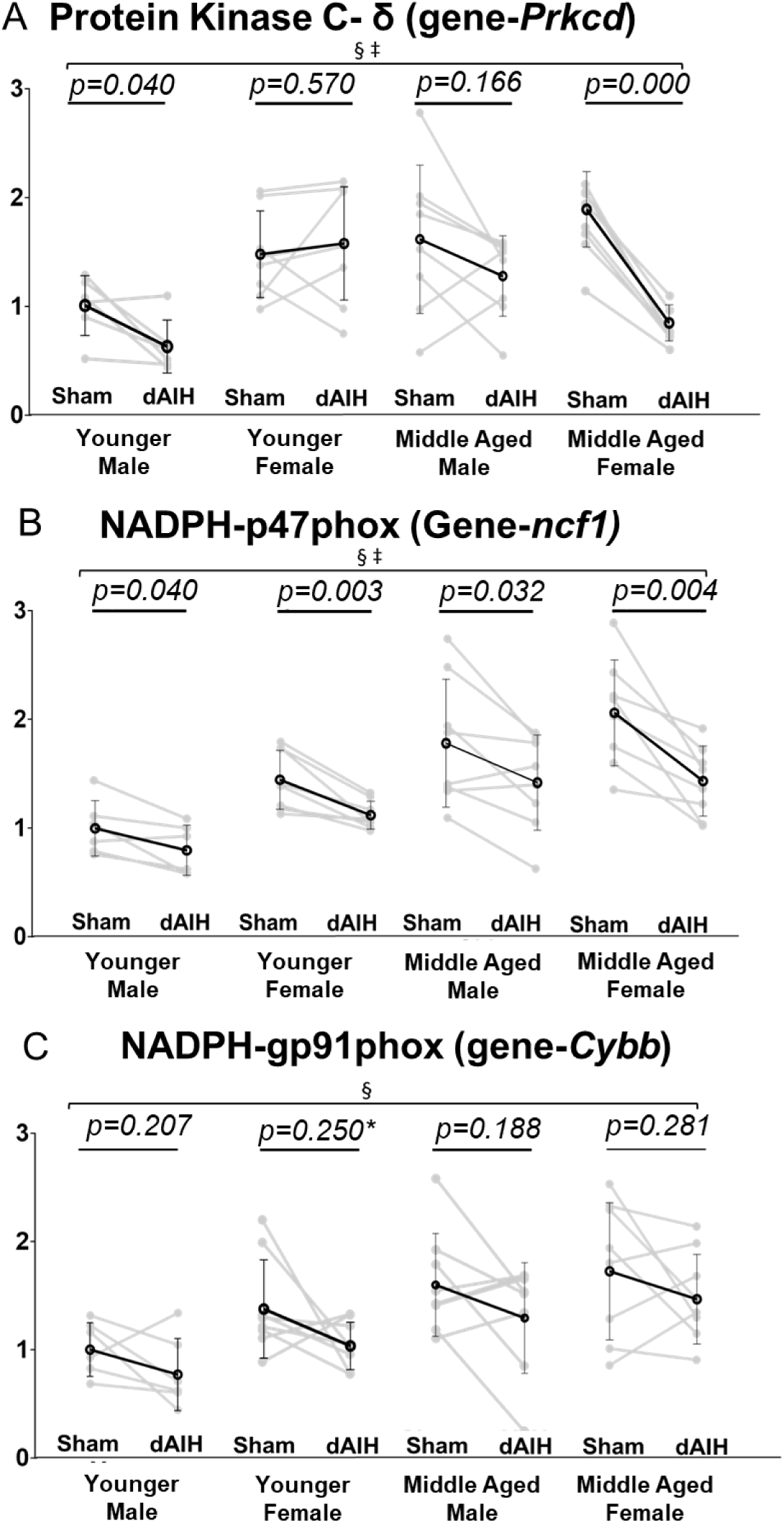
Effect of age, sex and daily AIH (dAIH) on genes associated with Q to S cross talk inhibition–PKCδ (*Prkcd*), NAPDH-p47 (*Ncf1*), and NAPDH-gp91 (*Cybb*). A significant effect of sex was observed in *Prkcd* expression, with greater expression in females. dAIH significantly reduced *Prkcd* only in middle aged females (Panel A). Significant age and sex effects were observed in *ncf1* expression, with greater expression in females. dAIH significantly reduced *ncf1* in young and middle aged females (Panel B). A significant increase in *Cybb* expression was observed with age, but no effect of sex or dAIH was observed (Panel C). §-Significant effect of age; ‡-Significant effect of sex; *-Non-parametric tested used due to lack of normal distribution.

### Effect of dAIH on S-to-Q crosstalk molecule gene expression

No significant changes in dAIH effects on S-to-Q cross-talk molecule gene expression were observed in younger males. However, a significant reduction in *Mapk14 was detected in* younger (Sham=1.68±0.45; dAIH=1.24±0.31; p=0.014) and middle-aged females (Sham=1.38±0.30; dAIH=1.07±0.26; p=0.011). A significant reduction in *Mapk11* expression was also observed in middle-aged males (Sham= 1.30±0.36; dAIH= 0.91±0.28; *p=0.003*), and in younger females (Sham=1.68±0.45; dAIH=1.24±0.31; p=0.007). *Pde4b* gene expression was reduced only in younger females (Sham=1.42±0.19; dAIH=0.94±0.14; p=0.003). Middle-aged females exhibited a significant reduction in *Prkaa1* (Sham=1.50±0.37; dAIH=1.15±0.11; p=0.014). No significant changes in *EPAC1* (**Figure 8**) or *Prkar1a* (**Figure 6**) were detected in any dAIH-exposed group (See Table 3).

**Figure 6:**
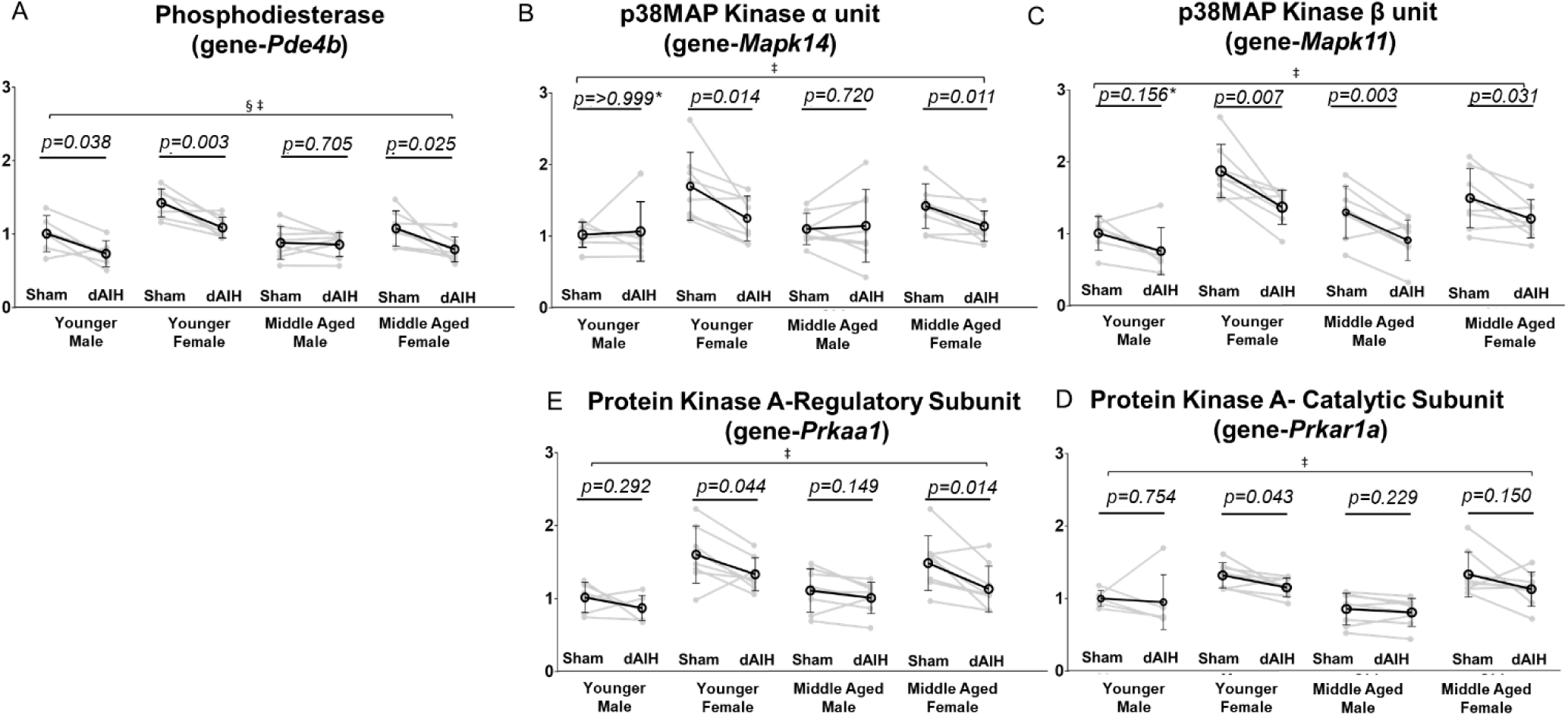
Effect of age, sex and daily AIH (dAIH) on genes associated with S to Q cross talk: phosphodiestease (*Pde4b*), p38Map kinase α (*Mapk14*), p38Map kinase β (*Mapk11*), PKA regulatory subunit (*Pkraa1*), and PKA catalytic subunit (*Prkar1a*). Females exhibited significantly reduced *Pde4b* gene expression with age, particularly in young females. dAIH significantly reduced *Pde4b* mRNA only in young females (Panel B). *Mapk14* and *Mapk11* were significantly greater in females, regardless of age (Panel C, D). dAIH significantly reduced *Mapk14* in young females (Panel C), whereas *Mapk11* was significantly reduced in younger females and middle aged males (Panel D). Significantly greater *Prkaa1* and *Prkar1a* expression were observed in females, regardless of age (Panel E, F). dAIH significantly decreased *Prkaa1* expression in middle aged females (Panel F), but not in any other group. §-Significant effect of age; ‡-Significant effect of sex; *-Non-parametric tested used due to lack of normal distribution.

### dAIH effects on *Cx3CL1* (fractalkine) expression

No significant change in *Cx3cl1* expression was observed in younger males (Sham= 0.96± 0.53; dAIH= 0.63±0.20; *0.156*) or females (Sham= 1.54±0.58; dAIH= 1.38±0.80; *p=0.326*), or middle-aged males (Sham= 1.40±0.35; dAIH= 1.52±0.21; *p=0.476*). However, *Cx3cl1* was was significantly reduced in middle-aged females (Sham= 1.87±0.35; dAIH= 0.85±0.18; *p=0.000*) (see **Figure 7** and Table 3).

**Figure 7:**
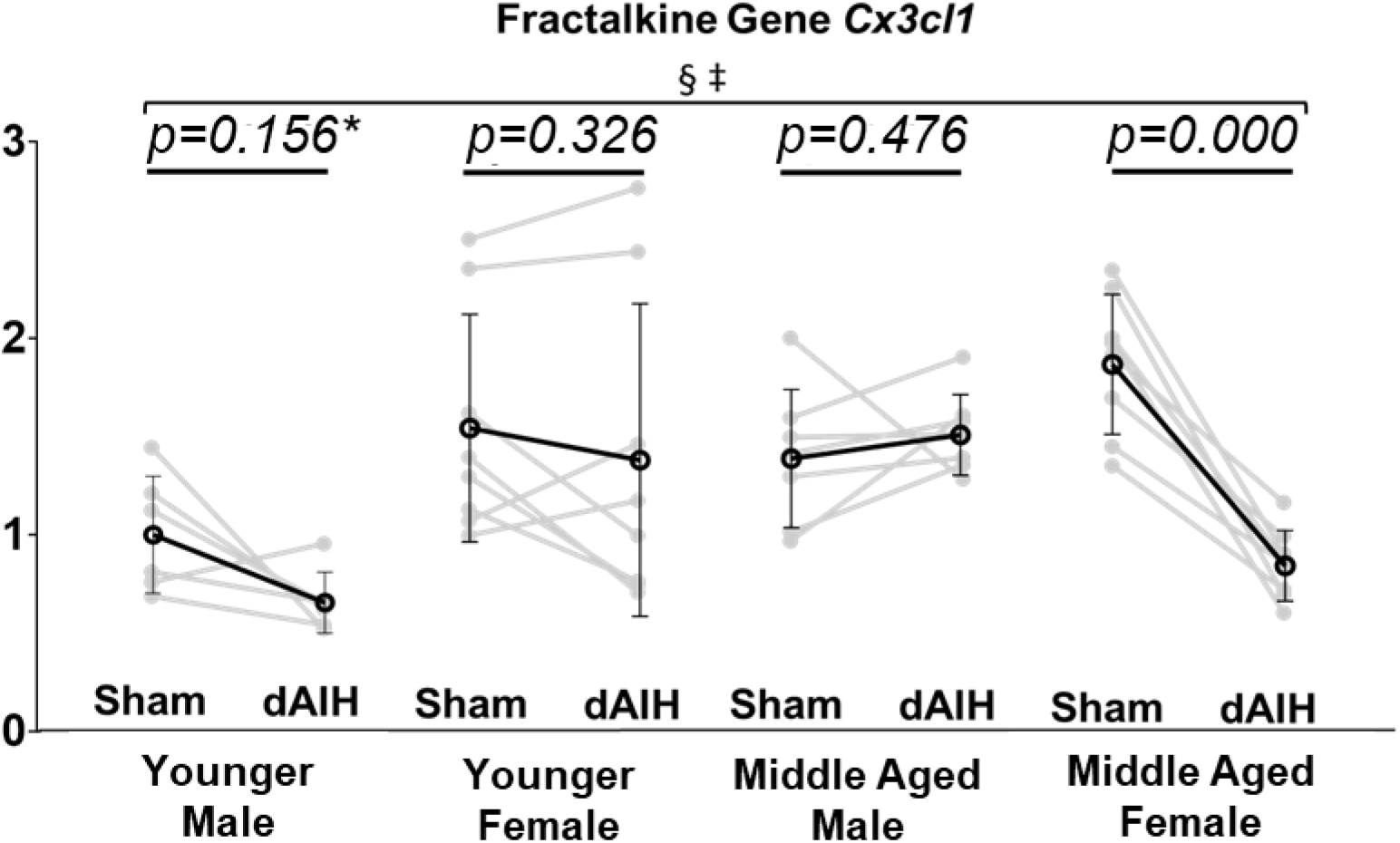
Effect of age, sex and daily AIH (dAIH) on neuron-microglia signaling molecule– Fractalkine (*Cx3cl1)*. Significant age and sex effects on *Cx3cl1* mRNA were observed; *Cx3cl1* mRNA higher in older rats, and in females regardless of age. dAIH significantly reduced *Cx3cl1* mRNA, but only in middle aged females. §-Significant effect of age; ‡-Significant effect of sex; *-Non-parametric tested used due to lack of normal distribution.

### dAIH effects on *Bdnf* gene expression

No significant change in *Bdnf* gene expression was observed in younger males (Sham= 1.00±0.33; dAIH= 0.70±0.39; *p=0.239*) or females (Sham= 1.12±0.16; dAIH= 1.27±0.40; *p=0.400*), nor were effects apparent in middle-aged males (Sham= 0.77±0.34; dAIH= 0.71±0.28; *p=0.641*) or females (Sham= 1.07±0.20; dAIH= 0.97±0.23; *p=0.365*) (see **Figure 8** and Table 3).

**Figure 8:**
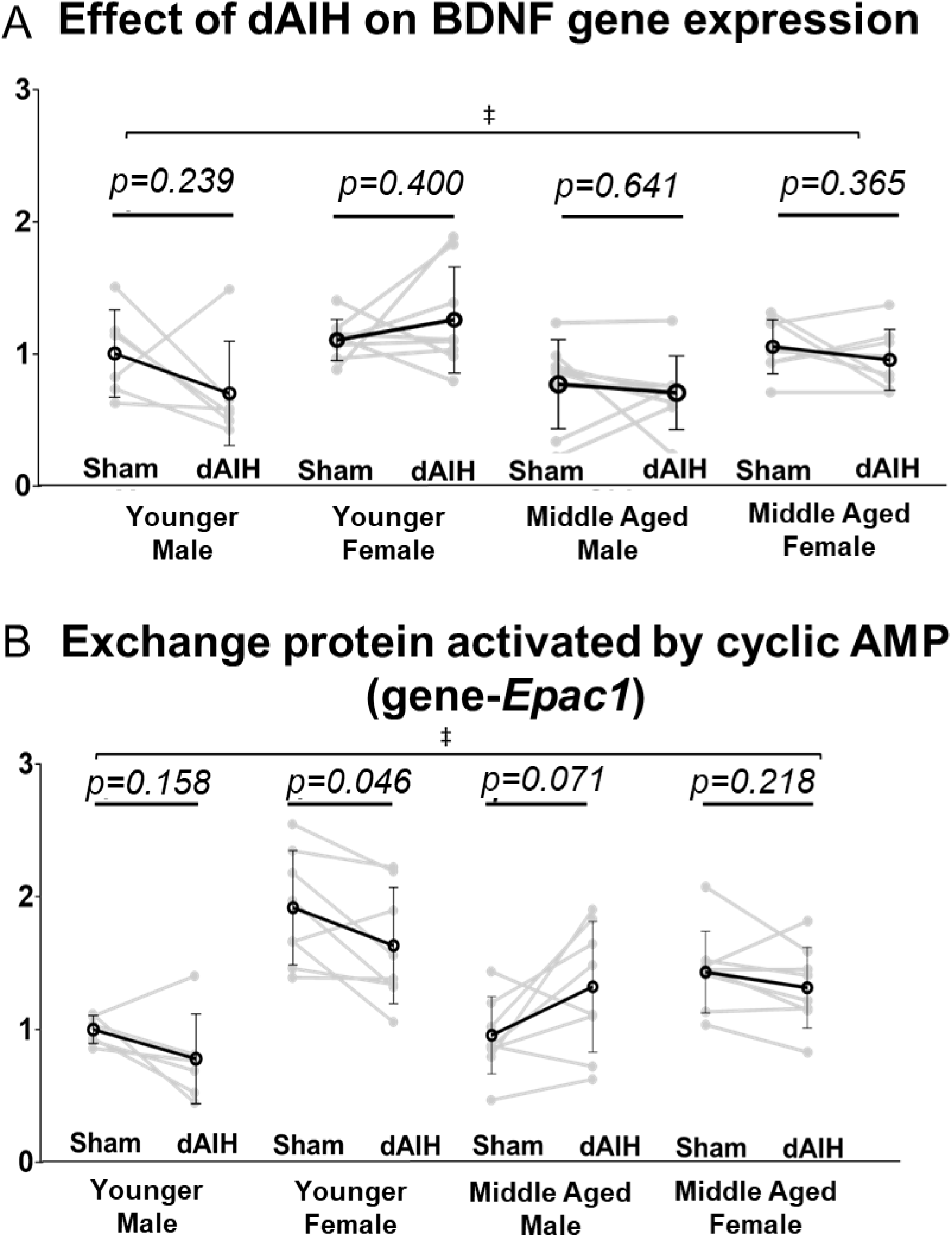
Age, sex and daily AIH (dAIH) effects on the pro-plasticity molecule, Brain derived neurotrophic factor (*Bdnf*) and Exchange protein activated by cyclic AMP (*Epac1*) mRNA. *Bdnf* gene expression was significantly greater in females regardless of age. dAIH had no effect on *Bdnf* expression in any group. EPAC (*Epac1*), an S pathway driver also activated by cAMP is also shown. Significantly greater *Epac1* expression was observed in females, regardless of age, but no age or dAIH effects were observed (Panel B). §-Significant effect of age; ‡-Significant effect of sex.

### Correlation of serum estradiol with Q-S crosstalk molecules

In sham control rats, serum estradiol positively correlated with *Epac1* (R^2^=0.29, p=0.002), *Mapk14* (R^2^=0.31, p=0.001) (**Figure 8**), *Mapk11* (R^2^=0.20, p=0.014), and *Prkar1a* (R^2^=0.20 p=0.013) mRNA. No other molecules correlated significantly with serum estradiol levels (Table 4).

**Table 4:**
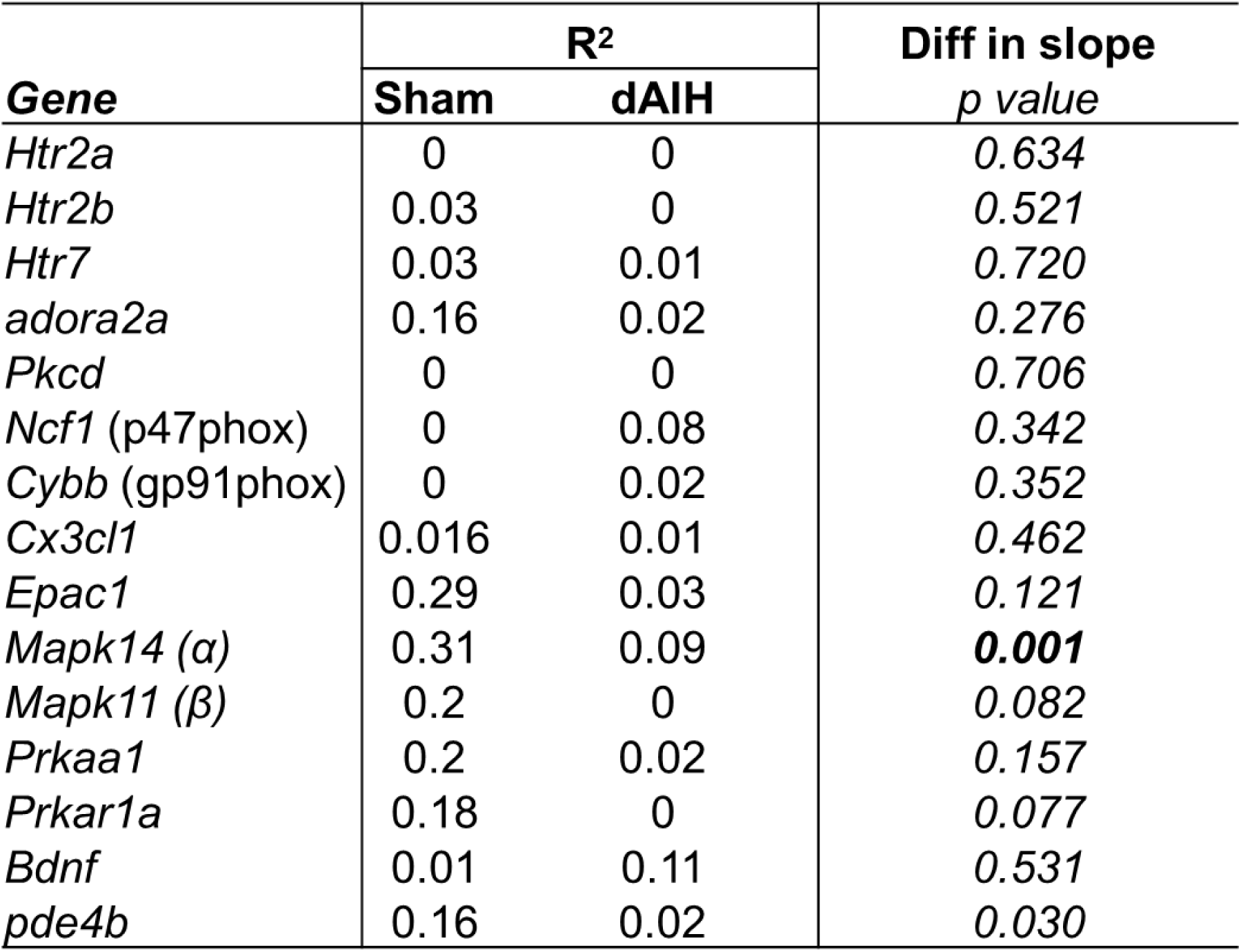
Serum estradiol level on daily acute intermittent hypoxia (dAIH) induced gene expression changes. Significance <0.01.

### Correlation of serum estradiol with dAIH effects

A significant correlation was observed between serum estradiol and dAIH-induced decrease in *MapK14* (Sham: R^2^=0.31; dAIH: R^2^=0.09, slope difference F=11.91, p=0.001; **Figure 9**). Serum estradiol was not significantly associated with dAIH-induced changes in Q or S pathway receptors (*Htr2a, Htr2b, Htr7*, and *Adora2a*), Q-to-S cross-talk (*Prkcd, Ncf1,* and *Cybb*) or S-to-Q cross-talk molecules (*Epac1, Prkaa1,* and *Prkar1a*), *Pde4b*, *Cx3cl1*, or *Bdnf* (Refer to Table 4).

**Figure 9:**
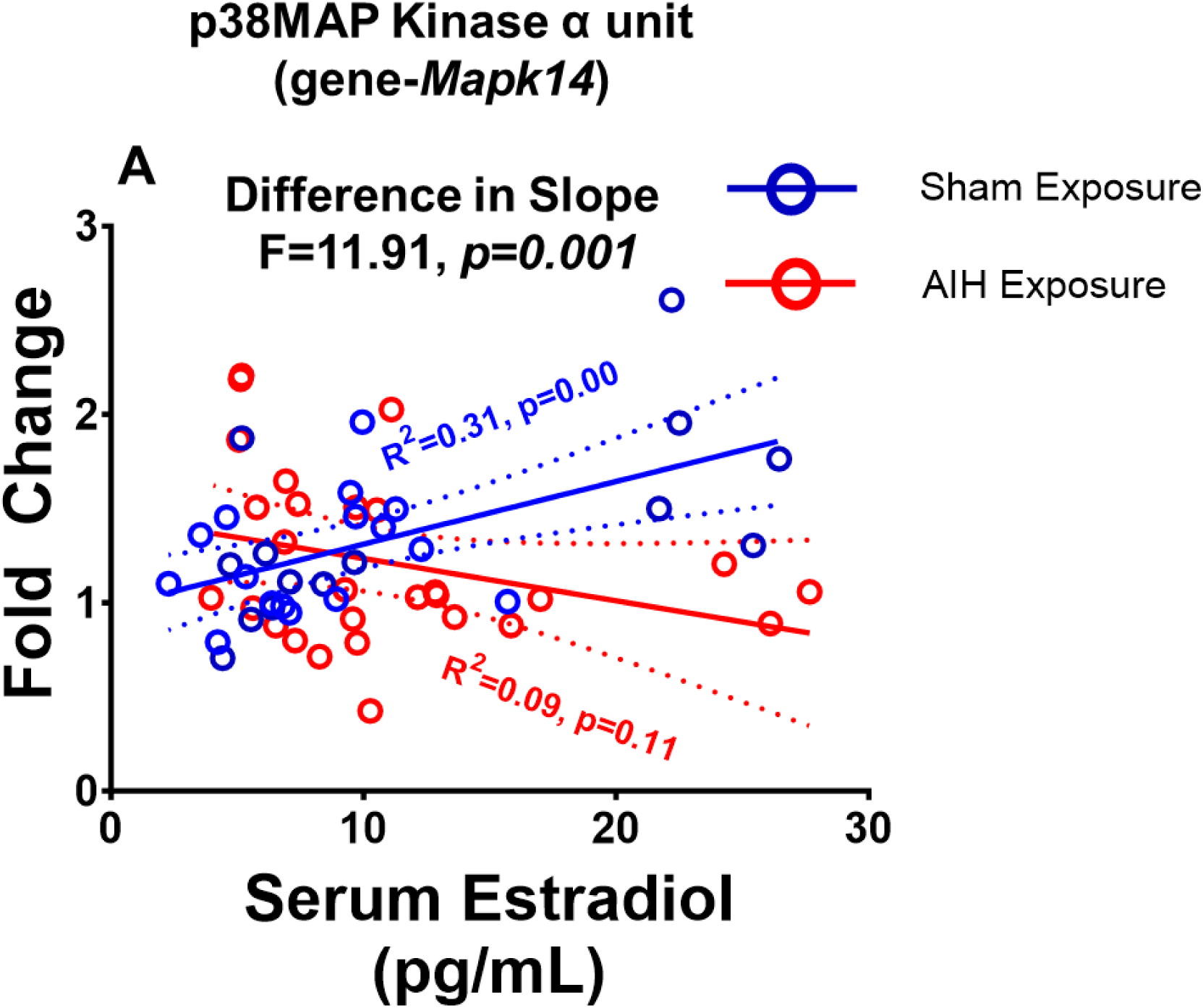
Correlation of serum estradiol with dAIH induced changes in p38 MAP kinase α unit (*Mapk14*) expression. A significant positive correlation was observed with increased serum estradiol in Sham rats. In rats exposed to dAIH, estradiol negatively correlated with *Mapk14*.

## DISCUSSION

We report important insights relevant to dAIH-induced metaplasticity in AIH-induced phrenic motor plasticity particularly concerning dAIH-induced changes in molecules known to regulate phrenic LTF, including molecules that mediate Q-S crosstalk inhibition. Further, we demonstrate that age, sex and circulating estrogen levels have an important impact on these changes (**Figure 10**). To summarize: 1) baseline plasticity-regulating genes vary in expression between young and middle-aged males and females; 2) dAIH-induced changes in Q or S pathway, and crosstalk molecules vary between young and middle-aged males *versus* females; 3) dAIH generally tends to decrease Q-S receptor and crosstalk molecules, particularly in middle-aged females; 4) to our surprise, dAIH had no effect on *Bdnf*; and 5) serum estradiol positively correlates with dAIH-induced decreases in Q-S crosstalk molecules, especially *Mapk14*. We discuss possible implications for age, sex and dAIH effects on phrenic motor plasticity, and limits of this study on ventral spinal homogenates.

**Figure 10:**
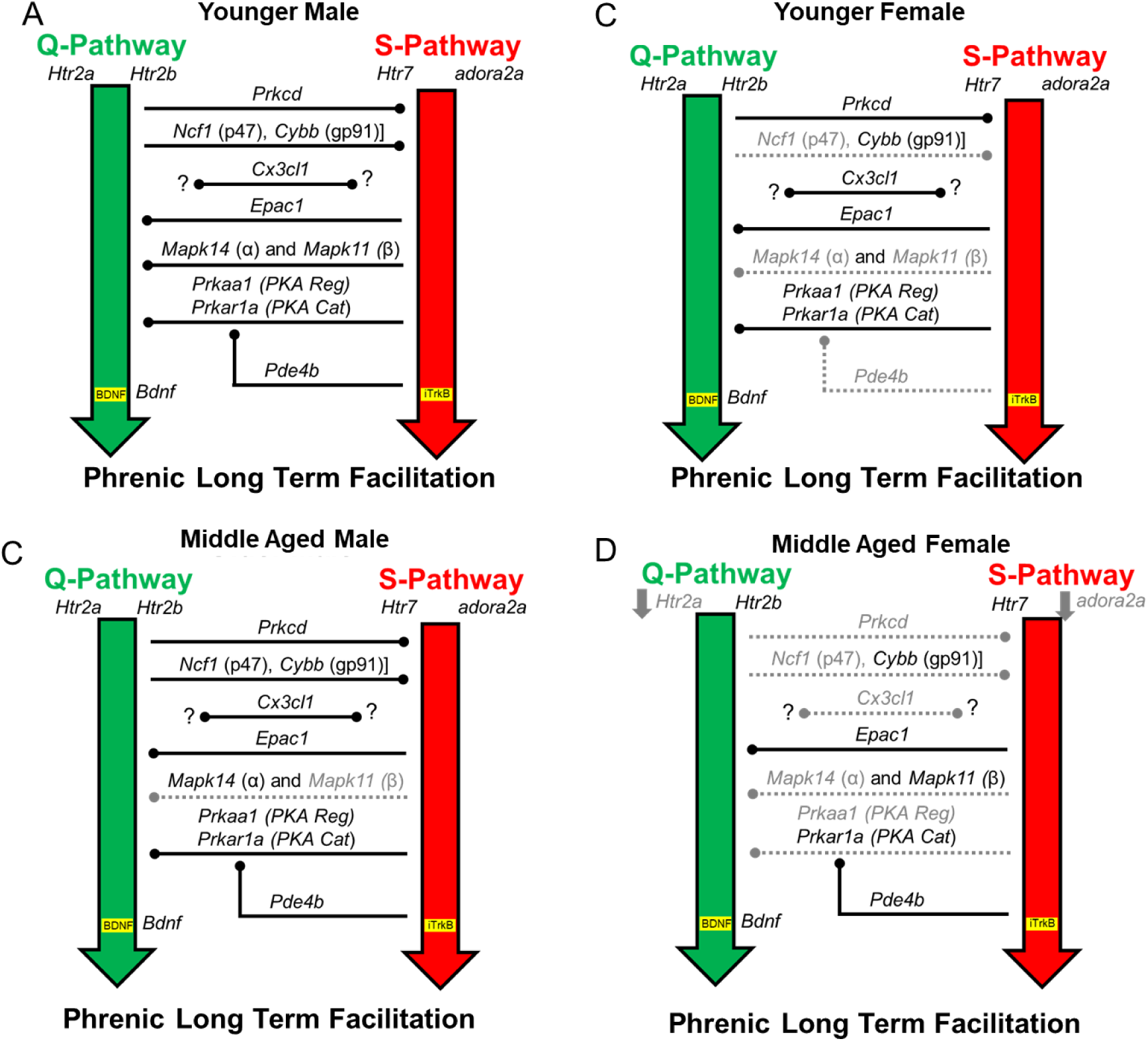
Conceptual diagram of dAIH effects on molecules engaged in Q-S cross talk inhibition in young (3 month) and middle aged (12 months) male and female rats. In young males, dAIH did not significantly change any crosstalk molecule investigated (Panel A). In young females dAIH significantly reduced Q to S cross talk molecules – NADPH-p47 (*ncf1*) and S to Q cross talk molecules – p38Map kinase α (*Mapk14*) and β (*Mapk11*). dAIH reduced phosphodiesterase (*Pde4b*) expression in young females, which could indirectly increase PKA or EPAC activity (Panel B). In middle aged males, dAIH had no significant effects on molecules involved in Q to S cross talk. However, S to Q cross talk molecule p38Map kinase β (*Mapk11*) expression decreased following dAIH (Panel C). In middle aged females, dAIH reduced Q to S cross talk molecules– PKC δ (Prkcd) and NADPH-p47 (Ncf1). S to Q cross talk molecules– PKA regulatory subunit (*Prkaa1*) and p38Map kinase α (*Mapk14*) were reduced following dAIH. dAIH also reduced 5HT2a (*Htr2a*) and A2a (*Adora2a*) expression in middle aged females (Panel D).

### Age-sex dimorphism in AIH induced phrenic LTF

Multiple studies indicate profound age dependent sexual dimorphism in phrenic LTF in rats. For instance, moderate AIH induced phrenic LTF decreases between young adult (3 months) to middle-aged (1 year) males, but increases with age in females (Behan et al., 2002; Behan et al., 2003; Dougherty et al., 2017; Zabka et al., 2001a, 2001b; Zabka et al., 2006). Since phrenic LTF also varies with estrus cycle (Zabka et al., 2001b), the impact of age, sex and estrus cycle on dAIH-induced metaplasticity must be considered. To date, nearly all studies of key Q and S pathway molecules were based on investigations in young adult male rats. Information concerning Q or S pathway molecules in middle-aged or geriatric males or females are scarce (but see: Marciante et al., 2023c; (Marciante & Mitchell, 2023). Thus, one major goal of this study was to understand maturation effects (young to middle-aged) on key Q versus S pathway molecules in male and female rats, their association with serum estradiol levels, and their impact on dAIH-enhanced phrenic long-term facilitation.

### Age-dependent sexual dimorphism and Q pathway gene expression

With advancing age in both sexes, there was an increase in Q pathway linked 5HT2b (*Htr2b*), PKCδ (*Prkcd*), NADPH (p47 phox (*Ncf1*); and gp91 (*Cybb*)) expression, without corresponding change in 5HT2a receptor expression. 5HT2b receptor-induced phrenic motor facilitation requires NADPH oxidase (NOX) activity (MacFarlane et al., 2011); NOX is also necessary for 5-HT2B crosstalk inhibition of the S pathway (Perim et al., 2018). Since p47 phox (*Ncf1*) and gp91 (*Cybb*) expression increase with age, we predict a corresponding increase in the potential for NOX-linked reactive oxygen species formation, although this would require mRNA translation to protein and enzyme activation. Reactive oxygen species inhibit adenyl cyclase activity and cAMP formation, providing a potential mechanism through which ROS undermine the S pathway (Haenen et al., 1990) and/or facilitate 5-HT2b induced Q pathway to phrenic motor facilitation (MacFarlane et al., 2011).

The protein kinase C δ isoform (PKCδ) constrains 5-HT7 receptor induced phrenic motor facilitation (Perim et al., 2019). PKCδ is a downstream target of estrogen receptors, and regulates mitochondrial respiration by controlling ROS formation (Evola et al., 2018; Jeong & Shin, 2012; Rex et al., 2010). PKCδ has a catalytic domain and a highly reactive regulatory domain, C1B, which reacts to diacylglycerols to inhibit adenyl cyclase and cAMP formation (Lai et al., 1997; Shanmugasundararaj et al., 2012). Thus, the age-dependent increase in PKCδ (*Prkcd*) expression, with yet higher expression in females, has important implications for Q to S pathway inhibition. Although PKC expression is typically assumed to decrease with age due to epigenetic modifications, PKCδ is an exception (Pascale et al., 1996; Patterson et al., 2010). Indeed, our results are consistent with one prior report demonstrating that PKCδ is not reduced with aging (Pascale et al., 1996). Since, PKCδ activity constrains the S pathway (Perim et al., 2019), increased i*Prkcd* gene expression might increase AIH induced phrenic LTF in middle-aged females by liberating the Q pathway from this inhibitory constraint.

### Age-dependent sexual dimorphism and S pathway gene expression

During severe hypoxia, glia-neuron interactions increase extracellular adenosine levels, activate A2a receptors (Acton & Miles, 2015; Boison et al., 2010), increase p38MAP kinase levels (Huxtable et al., 2015) and elevate chemokines unique to neurons (*e.g.,* fractalkine; *Cx3cl1*)(Arnoux & Audinat, 2015; Pawelec et al., 2020). Each of these molecules alone is sufficient to induce phrenic motor facilitation via the S pathway (Dale-Nagle et al., 2010; Golder et al., 2008; Hoffman et al., 2010; Hoffman et al., 2012; Nichols et al., 2012) (Fields & Mitchell, 2017; Hoffman & Mitchell, 2013). Our finding of an age associated increase in A2a receptor (*Adora2a)* expression supports a prior study (Marciante & Mitchell, 2023). Age-related increases in extracellular adenosine concentration (Castillo et al., 2009; Sebastião et al., 2000; Stockwell et al., 2017) increase A2a receptor activation and constrain the Q pathway to phrenic motor facilitation (Marciante & Mitchell, 2023). Downstream A2a receptor activation includes p38 MAP kinase activation in response to increased reactive oxygen species formation (Kralova et al., 2008). Although no age effect was observed in p38 MAP kinase genes in males, greater *MapK14* and *MapK11* expression with age in females and suggest potential differences in p38-dependent crosstalk inhibition between sexes.

Fractalkine (*Cx3cl1*) is a chemokine that, in the CNS, is uniquely expressed in neurons; its receptor is expressed exclusively by microglia (Pawelec et al., 2020). Recent studies indicate that phrenic motor neurons release Fractalkine during severe hypoxia, activating microglial Fractalkine receptors and triggering microglia-dependent adenosine accumulation necessary for expression of severe AIH-induced phrenic LTF (*i.e*., the S pathway); lesser adenosine accumulation during moderate hypoxia is only sufficient to constrain, but not replace serotonin-dominant phrenic LTF. Thus, the age associated Fractalkine mRNA increase found here (**Figure 7**) could upregulate neuron-microglia communication, increasing suppression or replacing Q pathway driven phrenic LTF. Overall, greater *Adora2a and Cx3cl1* expression in younger and middle-aged females suggests complex mechanisms involving age- and sex-dependent differences in neuron-glia interactions that reduce phrenic LTF in young adult females—the group with the highest expression level in all groups tested (Zabka et al., 2001b) and the lowest level of phrenic LTF (Zabka et al., 2001a, 2001b).

During severe hypoxia, increased intracellular levels of cAMP drives S pathway dependent phrenic motor facilitation *via* activation exchange protein directly activated by cAMP (EPAC) (Fields & Mitchell, 2017; Fields et al., 2015). In contrast, cAMP-triggered PKA signaling mediates cross-talk inhibition from the S to the Q pathway (Fields & Mitchell, 2017; Hoffman & Mitchell, 2013). Increased cAMP binds to PKA regulatory subunits (PKA Reg), inducing PKA conformational changes that free the catalytic C subunit (PKA C) to engage in enzymatic activity (Shaikh et al., 2012). Once activated, PKA C subunit phosphorylates cytosolic p47phox subunit, (Kandel, 2012) inhibiting NOX complex activation and reactive oxygen species formation (Bengis-Garber & Gruener, 1996; Kim et al., 2007; Nogueira-Machado et al., 2003). Diminished reactive oxygen species would reduce Q-S cross talk inhibition associated with 5HT2B receptor activation (Perim et al., 2018). During moderate hypoxia, marginal increases in cAMP are sufficient to activate protein kinase A (PKA), but not EPAC, the downstream effector of S pathway drive phrenic motor facilitation (Fields & Mitchell, 2017; Fields et al., 2015). Although no changes in PKA Reg (*Pkaa1*), PKA C (*Pkar1a*) or EPAC (*Epac1*) expression were observed with age, greater *Pkaa1, Pkar1a and Epac1* expression was observed in females regardless of age, suggesting the possibility of increased PKA-dependent Q pathway inhibition in females. Conversely, the long isoform of phosphodiesterase-4 (*Pde4b*), a potent inhibitor of PKA activity, likely plays an important role in balancing Q versus S cross-talk interactions (Fertig & Baillie, 2018). Thus, the decrease in *Pde4b* expression with age is consistent with reduced PKA-dependent S-to-Q cross talk inhibition. On the other hand, greater *Pde4b* expression in younger females suggests the potential for compensatory mechanisms, possibly offsetting PKA-mediated S to Q cross-talk inhibition. The overall impact of these changes on plasticity will depend on their collective balance, which is age and biological sex.

### dAIH effects and phrenic motor facilitation

Preconditioning with repetitive AIH enhances AIH-induced phrenic LTF (MacFarlane et al., 2018; Wilkerson & Mitchell, 2009) and amplifies short and long latency evoked phrenic responses from the ventrolateral funiculus (Perim, Sunshine, et al., 2021). Enhanced phrenic LTF is thought to arise from increased glutamatergic synaptic strength, possibly due to heightened NMDA and/or AMPA currents elicited by intermittent serotonin receptor activation (Bocchiaro & Feldman, 2004; McGuire et al., 2008). Empirical evidence detailing molecular mechanisms of reinforcement of synaptic inputs to phrenic motor neurons due to dAIH remains uncertain.

### dAIH effects on Q and S pathway ligand/receptor gene expression

The lack of dAIH-induced *Htr2b, Htr7* or *Bdnf* expression changes in ventral cervical homogenates are inconsistent with immunohistochemical or ELISA studies of protein in intact rats, which demonstrated increased BDNF (Lovett-Barr et al., 2012; Wilkerson & Mitchell, 2009) and 5HT2a receptor protein within phrenic motor neurons following repetitive AIH (Satriotomo et al., 2012). Differences between mRNA (this study) versus protein levels could explain this apparent discrepancy. In studies using other, repetitive AIH protocols (10 (Satriotomo et al., 2012) or 4 weeks of AIH 3x per week (MacFarlane et al., 2018); or 28 days of dAIH (Ciesla et al., 2021), serotonergic innervation within the phrenic motor nucleus is minimally affected. However, our results support observations reported by MacFarlane et al (MacFarlane et al., 2018), who found no changes in *Bdnf, Htr2b* or *Htr7* mRNA in ventral cervical homogenates following 4 weeks of repetitive AIH (3x per week) in male rats (MacFarlane et al., 2018). In contrast with those studies on males, *Htr2a* and *Adora2a* mRNA decreased in middle-aged females dAIH after dAIH of 14 days. Mechanisms and implication of reduced *Htr2a* and *Adora2a* mRNA are uncertain, but they were independent of serum estradiol levels in females (see below), but were associated with reduced Fractalkine mRNA and, possibly, adenosine levels.

### dAIH effects on Q-to-S crosstalk molecules

No significant dAIH effects on Q-to-S cross-talk molecules– PKCδ (*Prkcd*), NADPH-associated p47 phox (*Ncf1*) or gp91 (*Cybb*) mRNA were observed in younger or middle-aged males. However, the significant decrease in p47 phox (*Ncf1*) in young and middle-aged females suggests sexual dimorphism in Q-S pathway inhibition. Further, decreased PKCδ (Prkcd) in middle-aged females may support dAIH-enhanced plasticity in middle-aged females (Zabka et al., 2001b).

### dAIH effects on S-to-Q cross talk molecules

dAIH decreased key S-to-Q cross-talk inhibitory molecules including p38 MAP kinases–*Mapk14* (α) in younger and middle-aged females and *Mapk11* (β) in younger females and middle-aged males. Decreases in p38MAP kinase gene expression suggest the possibility of decreased neuroinflammation and, thus, Q pathway inhibition (Agosto-Marlin et al., 2018). Counterintuitively, decreased *Pde4b* expression in younger females is consistent with increased PKA activation and (possibly) greater S-Q cross talk inhibition; if the change in *Pde4b* is sufficient, it may activate EPAC sufficiently to drive the S pathway in response to AIH, consistent with the apparent contribution of the S pathway to enhanced phrenic LTF in rats exposed to repetitive AIH (3x per week, four weeks (MacFarlane et al., 2018; Perim et al., 2020); or daily for 14 days, (Perim, Sunshine, et al., 2021). The functional impact of dAIH-reduced *Pde4b* expression is of considerable interest but is unknown at this time. In middle-aged females, decreased PKA Reg (*Prkaa1)* and Fractalkine *(Cx3cl1)* expression could contribute to enhanced phrenic LTF (Zabka et al., 2001a, 2001b). Lack of *Epac1* or *Prkar1a* expression changes in any dAIH-exposed group is inconsistent with a role in age and sex effects on dAIH induced metaplasticity.

### Correlation of serum estradiol with dAIH-altered gene expression

A growing body of evidence demonstrates estrus cycle and female sex hormone effects on AIH-induced phrenic motor plasticity (Behan et al., 2002; Lurk et al., 2022; McIntosh & Dougherty, 2019). In females, AIH induced ventilatory facilitation is detectable only during proestrus, when the systemic estrogen levels are high (Dougherty et al., 2017; McIntosh & Dougherty, 2019). Similarly, phrenic LTF in middle-aged female rats (Zabka et al., 2003) requires a surge in systemic estradiol as rats transition to reproductive senescence (Frick, 2009). In males, gonadectomy abolishes phrenic LTF in young adult males, an effect restored by supplemental testosterone; however, this restoration of phrenic LTF requires conversion of testosterone to estrogen in the CNS via aromatase activity (Zabka et al., 2006). Thus, CNS estrogen likely impacts key molecules regulating the Q and/or S pathways and their cross talk inhibition.

We did not study estrous cycle effects in the present study, but rather correlated dAIH-induced changes with serum estradiol levels. In these naïve rats, serum estradiol positively correlated with *Epac1*, *Mapk14*, *Mapk11*, and *Prkar1a* mRNA. Although dAIH influenced mRNA for each of these molecules, a significant correlation between serum estradiol and dAIH-induced changes was observed only for *MapK14* mRNA. Decreased p38 MAP kinase subunits, specifically *Mapk14* (α), after dAIH suggests pro-plasticity and neuroprotective dAIH effects may be under the influence of circulating estradiol.

## LIMITATIONS

Ventral cervical (C3-C6) homogenates contain many cell types including neurons, microglia, astrocytes and oligodendrocytes. Thus, tissue homogenates from the ventral cervical spinal cord is unlikely to reflect gene expression within motoneurons *per se*. Further, estradiol concentrations in the ventral cervical spinal cord may differ from serum or even brain estradiol levels. Although progesterone regulates ventilation (Behan et al., 2003), the role of progesterone in AIH induced respiratory motor plasticity is difficult to study due to truncated duration of estrus cycle in rats (∼4 days) *versus* the human menstrual cycle (∼28 days) (McIntosh & Dougherty, 2019). We did not use comprehensive gene expression analytics involving RNAseq or Quantseq, but rather adopted the strategy of analyzing specific mRNAs from molecules known to regulate AIH-induced phrenic motor plasticity. Using this approach, we have a good idea of how changes in those molecules would impact plasticity, but may miss other molecules impacted by dAIH, age and/or sex (without knowledge of links to phrenic motor plasticity). Future studies using this discovery-based (versus hypothesis testing) approach to identify new molecules—although follow up studies will be needed to determine if they relate to any mechanism of phrenic motor plasticity.

## CONCLUSION

Based on this extensive study of age–dependent sexual dimorphism in daily AIH responses of molecules known to regulate phrenic motor plasticity, we conclude that daily AIH downregulates Q-S crosstalk molecules in an age and sex dependent manner, with the greatest downregulation in middle-aged females—when plasticity is at its peak (Zabka et al., 2001b). Based on correlations of estradiol with dAIH effects, we speculate that these variables regulate AIH-induced phrenic LTF at least in part by suppressing p38MAP kinase activity. This study begins to address an important question, but leaves open key questions such as: 1) is there a causal relationship between estradiol and the pro-plasticity molecules? 2) What cell types are changing expression of these molecules in response to daily AIH and/or age/sex; and 3) how do changes in these molecules impact plasticity? The novel findings presented here (and answers to the above questions) will advance our understanding of phrenic LTF, and inform translational research concerning the therapeutic potential of daily AIH to treat individuals of different ages and sex to improve breathing ability.

